# *ParaDISM*: Precise mapping of short reads to genes with highly homologous regions

**DOI:** 10.64898/2026.05.19.726275

**Authors:** Dimosthenis Tzimotoudis, Rosienne Farrugia, Jeremy Zammit, Maria Cini Masini, Alessandro Balestrucci, Francesca Borg Carbott, Stephanie Bezzina Wettinger, Panagiotis Alexiou, Michał Aleksander Ciach

## Abstract

**Background:** Genes with highly similar genomic copies (paralogs, tandem duplications and pseudogenes) pose a major challenge for Short-Read High Throughput Sequencing (srHTS). High sequence similarity makes it difficult to unambiguously identify the sequences of origin of short reads. This results in misalignment artifacts which can propagate through bioinformatic pipelines and increase error rates in variant calling.

**Results:** We present *ParaDISM*, a pipeline that refines standard alignments to improve read placement and reduce misalignment-driven false variant calls in highly homologous sequences. *ParaDISM* assigns a read/read pair to a sequence only when supported by unambiguous sequence-specific evidence by using a multiple sequence alignment of reference sequences to identify disambiguating positions. An optional iterative refinement procedure calls variants from confidently assigned reads, updates the reference sequences, and processes remaining non-assigned reads.

We evaluated the performance of *ParaDISM* both in terms of read alignment and the resulting short variant calls using extensive computational simulation experiments and the Genome in a Bottle HG002 benchmark. We applied *ParaDISM* to reanalyze two case studies: five public tumour exomes at the *GNAQ/GNAQP1* locus, and 18 short-read sequencing datasets of patients diagnosed with Autosomal Dominant Polycystic Kidney Disease (16 exomes and 2 panel sequencing datasets). Compared to the standard aligners (bowtie2, bwa-mem and minimap2), *ParaDISM* reduced the number of misalignment artifacts and false variant calls, resulting in an increased specificity and precision of the results.

**Conclusions:** *ParaDISM* improves the precision of read placement and single-nucleotide variant calling in highly homologous reference sequences. By reducing the number of false variant calls caused by misalignment artifacts, *ParaDISM* provides a stronger level of evidence for the called variants compared to currently available approaches. The pipeline is open source and available under the MIT license at github.com/BioGeMT/ParaDISM.

## Background

Short-read High Throughput Sequencing (srHTS) has transformed biomedical research and clinical practice by offering cheap, fast, and massive sequencing capabilities [1,2]. This technology enabled accurate and accessible detection of inherited (germline) variants for diagnosis of genetic disorders, as well as detection and analysis of somatic mutations in cancer for a better treatment and prognosis. However, many medically relevant genes remain either difficult or currently impossible to analyze with srHTS [3–5].

One of the major groups of problematic genes are ones that are highly similar to other regions of the genome, such as paralogs (genes originating from an ancestral duplication during the course of evolution), pseudogenes (generally non-functional copies of genes, often former paralogs), tandem duplications, and other highly repetitive regions. High sequence similarity (referred to as *sequence homology*) makes placement of short reads, and subsequent variant calling, inaccurate [6–11]. This is exacerbated by the design of many alignment algorithms, which either look for the first placement that is “good enough” rather than optimal, or discard reads which have multiple possible placements. As a consequence, misaligned reads contribute to errors in variant calling, resulting in false positives (incorrectly called variants) and false negatives (failure to detect a variant) [6,12].

A prime example of issues caused by homologous sequences is the diagnostics of the Autosomal Dominant Polycystic Kidney Disease (ADPKD). This is one of the most common inherited kidney diseases, with prevalence estimated at nearly 1 in 1000 people worldwide, predominantly caused by mutations in *PKD1* and/or *PKD2* [13,14]. *PKD1* has six pseudogenes with sequence homology up to 97% in some regions [15]. This poses a major challenge for accurate alignment of short reads, which significantly limits the diagnostic applicability of srHTS [16]. Therefore, even though whole-genome sequencing (WGS) can overcome the pseudogene problem in some cases [17,18], and research on optimizing srHTS is ongoing [19], it has not yet replaced long-range PCR followed by Sanger sequencing of nested amplicons as the gold standard [20]. This has important consequences for disease management, excluding disease in kidney donors, and efforts to design treatments such as the recent approaches based on microRNAs [21].

Another example of a possible misalignment artifact reported in the literature involves the *GNAQ* locus and its pseudogene *GNAQP1*. Recurrent *GNAQ* hotspot mutations (p.T96S and p.Y101X) were reported from short-read WES in natural killer/T cell lymphoma [22]. However, *GNAQ* shares a short region of very high sequence identity with *GNAQP1* that spans the hotspot sites. It was later demonstrated that common *GNAQP1* polymorphisms (rs3730150, rs3730148 and rs3730153) can cause short reads to be misaligned by standard aligners and mimic the reported *GNAQ* hotspot signature [6].

Some bioinformatic approaches to disambiguate between homologous sequences augment standard short-read pipelines with locus-specific steps, often focusing on a single gene or a limited set of duplicates [7,8,23]. For example, PMS2_vaR [23] improves short-read analysis of *PMS2* in the presence of the *PMS2CL* pseudogene by forcing alignments to the *PMS2* reference sequence and using gene-specific invariant positions to resolve reads from the problematic exon 11 region. Other approaches focus primarily on variant discovery and interpretation in paralogous regions at larger scales. For instance, Chameleolyser [24] reprocesses reads from paralogous loci by realigning them to masked reference sequences and, for indistinguishable paralogs, reports variants as having ambiguous locus assignment. Complementary benchmarking work provides nucleotide-resolution measures of short-read sequencing difficulty in exonic regions, e.g., the UNMET score, highlighting positions that are challenging for variant calling with conventional srHTS [25]. Due to the difficulty of accurate read placement, many diagnostic strategies for such genes rely heavily on experimental protocols and specialized laboratory workflows [26,27].

One approach is to use third generation long-read sequencing, which can span regions with less similarity and therefore allow for more accurate read placement [28–32]. However, using long reads is not free from challenges. Long reads do not always fully remove the possibility of misalignment artifacts on extensive regions of high similarity, necessitating the use of specialized software such as DuploMap or Paraphase to improve mapping and variant calling in duplicated regions [33,34]. For example, Paraphase separates haplotypes based on manually identified sites which differ between the reference homologous sequences, e.g. *SMN*c.840 to separate *SMN1* and *SMN2*.

A particular limitation of many of the aforementioned approaches is the focus on the genes directly responsible for disease. However, detecting and analyzing variants in homologous regions (such as pseudogenes or paralogs) not only can discover a false positive call in the gene of interest [6], but may also have clinical importance on its own. Pseudogenes have long been considered to lack function, but this view is becoming increasingly challenged [35]. Moreover, there have been documented gene conversions between *PKD1* and its pseudogenes [36]. This calls for an approach that can accurately map reads to a gene and all of its homologs to allow for a more accurate variant calling without extensive modifications of laboratory protocols.

In this work, we present *ParaDISM* (Paralog Disambiguating Mapper), a pipeline for paralog-aware placement of short reads in highly homologous genomic regions. It builds a multiple sequence alignment (MSA) of the reference homologous regions to identify disambiguating positions and aligns a read/read pair only when it finds unambiguous evidence for the sequence of origin; otherwise, the read is not assigned to any sequence and reported separately. The pipeline outputs sequence-specific alignments for inspection and downstream variant calling, and can be applied to panel, whole-exome (WES) and whole-genome (WGS) short-read sequencing data.

## Implementation

### The *ParaDISM* pipeline

An overview of the workflow is shown in Fig. 1. *ParaDISM* takes short-read sequencing data (single-end or paired-end FASTQ) and a user-generated FASTA MSA file of homologous sequences (all in a single orientation). The pipeline outputs a per-read (or per-read pair) gene assignment together with gene-specific FASTQ and BAM files for downstream inspection and analysis. Reads that cannot be placed unambiguously are labeled None and output in a separate FASTQ file.

**Fig.1:**
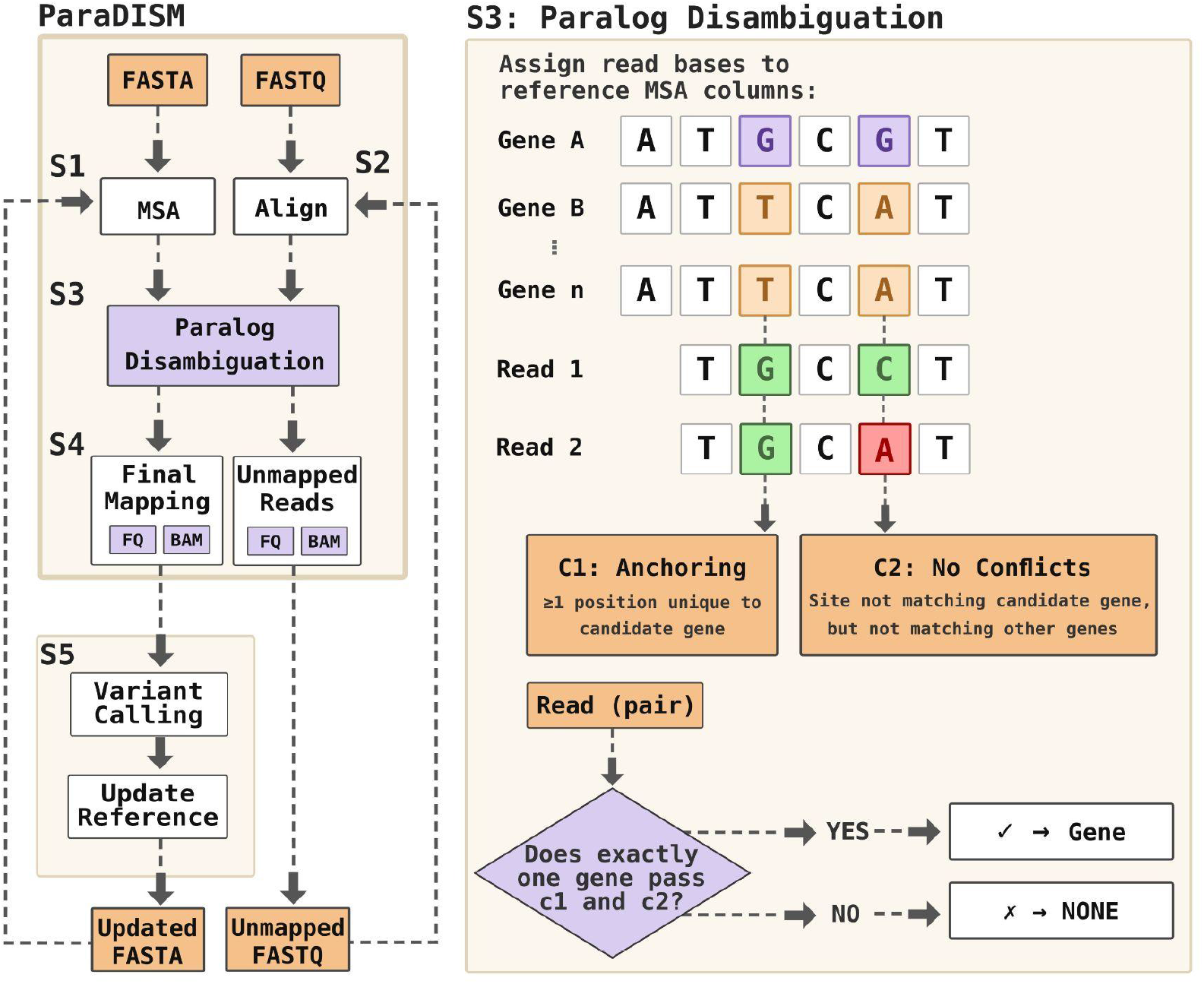
The ParaDISM pipeline architecture. The pipeline takes as input a FASTQ file with sequencing reads and a FASTA file with reference sequences of the homologous regions. Reads are assigned to a sequence only when there is unambiguous evidence that it is the true sequence of origin, expressed as two conditions (C1: Anchoring and C2: No Conflicts). In the example shown in the Figure, Read 1 passes both conditions, while Read 2 fails the C2 condition. The pipeline iteratively updates the reference sequences to identify sample-specific variants which may interfere with the alignment conditions or serve as additional anchoring sites. Reads which could not be aligned are reported separately for a manual inspection.

#### *Step 1 (S1 in Fig. 1)*: Multiple Sequence Alignment

The reference sequences are aligned using MAFFT [37] to produce a multiple sequence alignment (MSA). The MSA is used to define homology relationships between the nucleotides of all the homologous reference sequences. *ParaDISM* uses this relation to identify positions that can disambiguate between homologous sequences: MSA columns in which a given sequence has a nucleotide that is different from the homologous nucleotides of all other sequences.

#### *Step 2 (S2 in Fig. 1):* Initial Read Alignment

HTS dataset reads are aligned to the reference sequences using a standard aligner (with the current implementation supporting Bowtie2 [38], BWA-MEM2 [39], or minimap2 [40]). The user can specify a minimum alignment score threshold to filter low-quality mappings and suppress off-target alignments. This initial alignment provides candidate placements and read-to-reference coordinates for the disambiguation step.

#### *Step 3 (S3 in Fig. 1):* Paralog disambiguation

For each read (or read pair), *ParaDISM* assigns aligned read bases to MSA columns and evaluates each reference sequence as a candidate gene assignment. For a given candidate, *ParaDISM* evaluates two conditions:

- Condition 1: Anchoring position (S3 C1 in Fig.1) the read contains at least one base that matches the candidate sequence but does not match any other sequence at the homologous MSA column.
- Condition 2: No conflicting positions (S3 C2 in Fig.1): the read does not contain any base that mismatches the candidate sequence while matching another sequence at the homologous MSA column.

A read is assigned to a sequence if it is the unique sequence which passes both conditions. For paired-end data, evidence is combined across both reads of a paired read (either read can provide an anchoring position, but neither read can introduce a conflict). If no candidate passes, or if multiple candidates pass, the read (read pair) is labeled None, indicating it could not be assigned to any sequence.

#### *Step 4 (S4 in Fig.1):* **Per-gene realignment and output generation**

*ParaDISM* writes sequence-specific FASTQ files for reads assigned to each sequence and re-aligns those reads to the corresponding sequence (each sequence separately) to obtain within-gene coordinates. The resulting sorted and indexed per-sequence BAM files can be used for visualization and downstream variant calling. Reads not assigned to any sequence by the disambiguating algorithm are reported in a separate FASTQ file and mapped to all reference sequences with a standard mapper to allow for a manual inspection of variants possibly existing in the genome but not supported by reads with unambiguous placement.

#### *Step 5 (S5 in Fig. 1):* Iterative refinement (optional)

In iterative mode, *ParaDISM* attempts to rescue reads initially labeled None by incorporating sample-specific variants into the reference sequences. Genetic variants present in a sample can disrupt anchoring positions and cause otherwise informative reads to be labeled None. *ParaDISM* calls variants from confidently mapped reads using FreeBayes (Garrison and Marth 2012), optionally applies quality filters (QUAL/DP/AF), updates the reference sequences with called variants, and re-runs the workflow on the remaining unmapped reads. Assignments from earlier iterations are preserved, and the procedure stops when no additional reads are reassigned or no new variants are found, or when a maximum number of iterations is reached.

### Validation on simulated datasets

Reference sequences of *PKD1* and its six pseudogenes, *PKD1P1-PKD1P6*, were downloaded from the Ensembl database (ENSG00000008710, ENSG00000244257, ENSG00000227827, ENSG00000183458, ENSG00000205746, ENSG00000254681,ENSG00000250251; human genome build GRCh38). Sequences of genes located on the negative strand *(PKD1, PKD1P4-6*) were reverse complemented. The sequences were used as a reference “genome” to simulate 1000 paired-end srHTS “panel” data sets with DWGSIM (Whole Genome Simulator for Next-Generation Sequencing; https://github.com/nh13/DWGSIM). Random seeds 1 to 1000 were used. Each data set consisted of 100,000 read pairs (read length 150 bp; insert size 350 ± 35 bp; per-base sequencing error rate 0.01; SNP rate 0.005; indel rate 0.0005; indel extension probability 0.5). Each data set was analyzed with *ParaDISM* with all three supported mappers (10 iterations; FreeBayes quality filters QUAL≥20, DP≥10, AF≥0.05, minimum alternate allele count 5).

### Genome in a Bottle (GIAB) HG002 benchmark

GIAB HG002 (NA24385) Illumina short reads and GIAB/NIST v4.2.1 truth set on GRCh38 were downloaded from the GIAB website (https://www.nist.gov/programs-projects/genome-bottle) [41]. Reads were merged across lanes. In addition to the full dataset, an approximate 10× whole-genome sequencing data set was generated by downsampling the number of read pairs to 62,746,019 (assuming 500 bp per read pair for 2×250 bp reads). *ParaDISM* was run with Bowtie2 in local mode for 10 iterations with *PKD1* and pseudogenes as the reference FASTA. A minimum Bowtie2 alignment score function of G,60,60 was used to suppress off-target alignments. For 250-bp reads, this setting corresponds to a minimum score of ∼391 (≈78% of a perfect-match score under Bowtie2’s default local scoring). Variants were called using Freebayes. Truth and callsets were normalized and decomposed into biallelic records (multi-allelic sites were split). Only biallelic SNPs were retained, and evaluation was restricted to GIAB benchmarkable regions. Final benchmark callsets were filtered using INFO/AO≥3, QUAL≥10, INFO/SAF≥1, and INFO/SAR≥1.

### GNAQ Case Study

Five whole-exome tumour datasets (SRR5602384, SRR5602389, SRR5602393, SRR5602414, SRR5602419) were downloaded from NCBI Sequence Read Archive. Reference GRCh38 sequences of *GNAQ* and its pseudogene *GNAQP1* were downloaded from the Ensembl database (ENSG00000156052, ENSG00000214077) and used as a two-sequence reference FASTA. *ParaDISM* was run with BWA-MEM2 for 10 total iterations using a minimum BWA-MEM2 alignment score threshold of 160. Variants were called with FreeBayes using the same filters as in the previous experiments. Read support was summarized using SAMtools mpileup [42] by computing per-base pileup counts at the *GNAQ* hotspot positions and at the three *GNAQP1* SNP positions reported by Lim et al. [6].

### ADPKD Case Study

Patients with ADPKD were recruited through the nephrology clinics (paediatric and adult) at Mater Dei Hospital, Malta, as part of the Malta NGS Project after written informed consent was obtained (ethics approval number FHS031/2014). The proband in each pedigree had typical ADPKD signs and symptoms as defined by the Barua and Pei criteria [43]. Blood or saliva were collected from 18 patients from 15 different families. Short-read high-throughput sequencing was carried out using either a custom target panel (n=3) or whole exome sequencing (n=15). Libraries for srHTS were prepared as per Agilent SureSelect^XT^ target enrichment protocol for paired-end Illumina sequencing. Sequencing was carried out on an Illumina HiSeq4000 platform. Shortlisted potentially causative variants were confirmed using a combination of long-range PCR, nested PCR and Sanger sequencing as previously described [20].

*ParaDISM* was run with Bowtie2 for 10 total iterations with a minimum Bowtie2 alignment score function of G,40,40. Variants were called using Freebayes with the same filters as in the previous experiments. To compare *ParaDISM* with a more standard, non pseudogene-aware approach, FASTQ files were analyzed with NextGENe v.2.4.2.3. In NextGENe, reads were trimmed or rejected when 3 or more bases had a Phred score of 20 or less. Cleaned raw reads were aligned to GRCh37 and variant calls were made (variant minimum coverage 20x, Phred score ⩾Q20). The resulting VCF files were compared between both approaches.

## Results and Discussion

### Precise placement of reads on highly homologous sequences

To evaluate the accuracy of *ParaDISM* alignment, we generated an extensive dataset of simulated srHTS experiments (where the gene of origin is known for each read pair), and used them to rigorously compare the performance of the proposed pipeline against standard solutions. We generated 1000 paired-end srHTS datasets that simulate panel sequencing of *PKD1* and its six pseudogenes. For each dataset, we compared the performance of *ParaDISM* against Bowtie2, BWA-MEM2, and minimap2 (short-read preset).

We used three performance metrics to compare the alignments: per-sequence sensitivity (the proportion of reads originating from a sequence that were successfully mapped to it), per-sequence specificity (the proportion of reads originating from other sequences that were not mapped to the analyzed sequence), and per-sequence precision (the proportion of reads mapped to the sequence that did originate from it). The first two metrics are standard approaches in medical literature and often used to evaluate approaches to variant calling, while the latter is used arguably less often. However, we note that specificity of read placement can be misleadingly high, because the data is inherently unbalanced: more reads originate from other sequences than from any given sequence. This is best illustrated by the following simplified example. Suppose we analyze 10 sequences, with equal coverage of each sequence. Even if the reads are assigned to the sequences completely at random, the expected specificity is equal to 90%. On the other hand, the expected precision in this case is equal to only 10%. Since specificity can be very high even for a completely random assignment, precision is more suitable for a rigorous evaluation of read placement.

On the simulated data, *ParaDISM* with 10 iterations improved the specificity and precision of read assignment compared to standard aligners (Fig. 2). Averaged across Bowtie2, BWA-MEM2, and minimap2, specificity increased from 0.97 to 0.99 and precision increased from 0.78 to 0.88. The improvement was consistent across all three aligners, which yielded very similar overall specificity (0.9687, 0.9688, and 0.9689, for BWA-MEM2, Bowtie2, and minimap2 respectively) and precision (0.7800, 0.7807, and 0.7799). Depending on the algorithm used for the initial alignment of reads in *ParaDISM*, the specificity increased to 0.9879, 0.9892, and 0.9898 for the three aligners respectively and precision increased to 0.8713, 0.8802, and 0.8839, indicating that the choice of aligner used by *ParaDISM* has a lower impact on the results than the paralog disambiguating step.

**Fig. 2:**
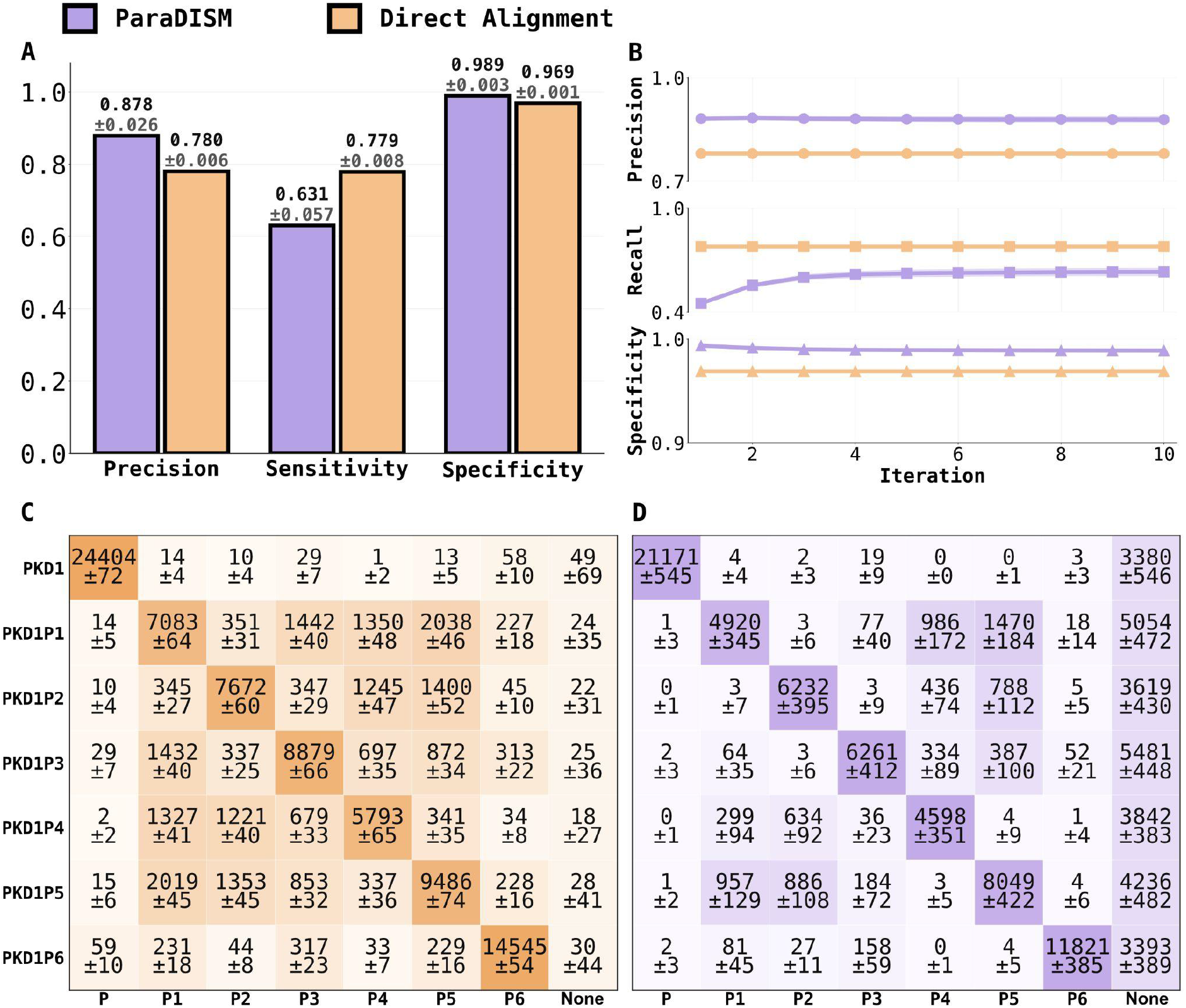
Read placement performance in PKD1/PKD1P simulations. (A) Overall precision, sensitivity, and specificity of ParaDISM versus standard aligners, averaged across 1,000 simulations and three aligners (Bowtie2, BWA-MEM2, minimap2), in a simulated PKD1 and pseudogene experiment. Numbers above the bars report mean ± standard deviation across the simulations. (B) Iteration-wise precision, sensitivity, and specificity during the iterative refinement procedure. The averaged metrics of the standard aligners are shown as a reference (constant for all iterations). (C) Read assignment matrix for the standard aligner. Rows indicate the true sequence of origin and columns indicate the assigned sequence. The None column denotes reads not assigned to any sequence. Cells report mean ± standard deviation read counts across 1,000 simulations. Misalignment artifacts are visible as off-diagonal cells of the matrices. (D) Read assignment matrix for ParaDISM after 10 iterations. While the decrease in misalignment artifacts, visible as lower counts in off-diagonal cells, comes at the cost of larger counts in the None column, the sensitivity remains at a sufficient level to align many reads to the analyzed genes.

Read assignment matrices (Fig. 2C,D) show that standard aligners produced numerous misalignment artifacts between homologous sequences, whereas *ParaDISM* reduced the artifacts by concentrating ambiguity in the None category. Depending on the alignment algorithm, *ParaDISM* assigned 27.7%-30.1% of read pairs to the None category, compared with <0.01%-0.6% for the standard aligners without paralog disambiguation. Accordingly, by design of the pipeline, the increased precision came at a cost of decreased sensitivity from 0.78 to 0.63 (Fig. 2A). The lower sensitivity reflects *ParaDISM*’s conservative handling of ambiguity: reads that cannot be placed unambiguously due to excessive sequence homology are labeled None instead of forcing an unreliable alignment. The iterative refinement procedure was essential for maintaining the decrease in sensitivity at an acceptable level (Fig. 2B). Accordingly, despite the lowered sensitivity, the number of reads aligned to the sequences was sufficient for downstream analyses, with 21171 read pairs aligned to PKD1 by *ParaDISM* compared to 24404 by the standard aligners on average.

*ParaDISM* has reduced misalignment artifacts between every pair of the analyzed sequences, as evidenced by lower numbers of off-diagonal assignments in read assignment matrices in Fig. 2 C,D. However, the prevalence of misalignment artifacts, and the relative performance in which *ParaDISM* resolved them, varied highly between the sequences. For both *ParaDISM* and the standard aligner, most misalignment artifacts occurred between the pseudogene sequences (e.g. out of reads originating from *PKD1P1*, 351 were placed on *PKD1P2* by the standard aligners, compared to only three by *ParaDISM*; 2038 were placed on *PKD1P5* by the standard aligners, compared to 1470 by *ParaDISM*). Most notably, the standard aligner placed 129 pseudogene-derived read pairs on *PKD1*, which corresponds to 38700 sequenced bases for 150 read length and represents a major risk for false positive SNV calls. In contrast, *ParaDISM* incorrectly placed only six reads on *PKD1*, significantly reducing the risk of these calls.

### Higher precision of variant calling on an established benchmark dataset

As a next evaluation of *ParaDISM*, we investigated whether the increased specificity and precision of read placement result in more credible variant calling. We used FreeBayes to call SNVs in *PKD1* and its pseudogenes using alignments produced by either *ParaDISM* with Bowtie2 as the initial mapper or Bowtie2 alone on the established Genome In A Bottle (GIAB) HG002 benchmark srHTS dataset with known variants [41].

We compared the performance of SNV calls using sensitivity and precision metrics. We note that the aforementioned issue with misleadingly high specificity is even more pronounced in SNV calls than in read placement. A caller which calls SNVs completely at random with probability 1% at each nucleotide has an expected specificity of 99% regardless of the length of the sequence or number of SNVs present. This is because specificity only measures how well a caller identifies sites *without* variants, and in those sites, this kind of a random caller would make an error with 1% probability. On the other hand, on a sequence length of 1000 bp with 1 SNV present, this kind of a random caller would have a precision of only 0.1%. Because of this, we decided not to compare the specificity of different approaches but use only the precision and sensitivity instead.

On the downsampled ∼10× dataset, *ParaDISM* produced 11 true positives and, notably, no false positives, resulting in a precision equal to 1.00 (Fig. 3 A). On the other hand, while FreeBayes applied to Bowtie2 alignments managed to find 29 true positives, it also produced eight false positives, resulting in a lower precision of 0.78. The reduction in false positive calls thanks to paralog disambiguation was most pronounced in pseudogenes. For example, on *PKD1P6, ParaDISM* resulted in zero false positives while Bowtie2 produced eight.

**Fig. 3:**
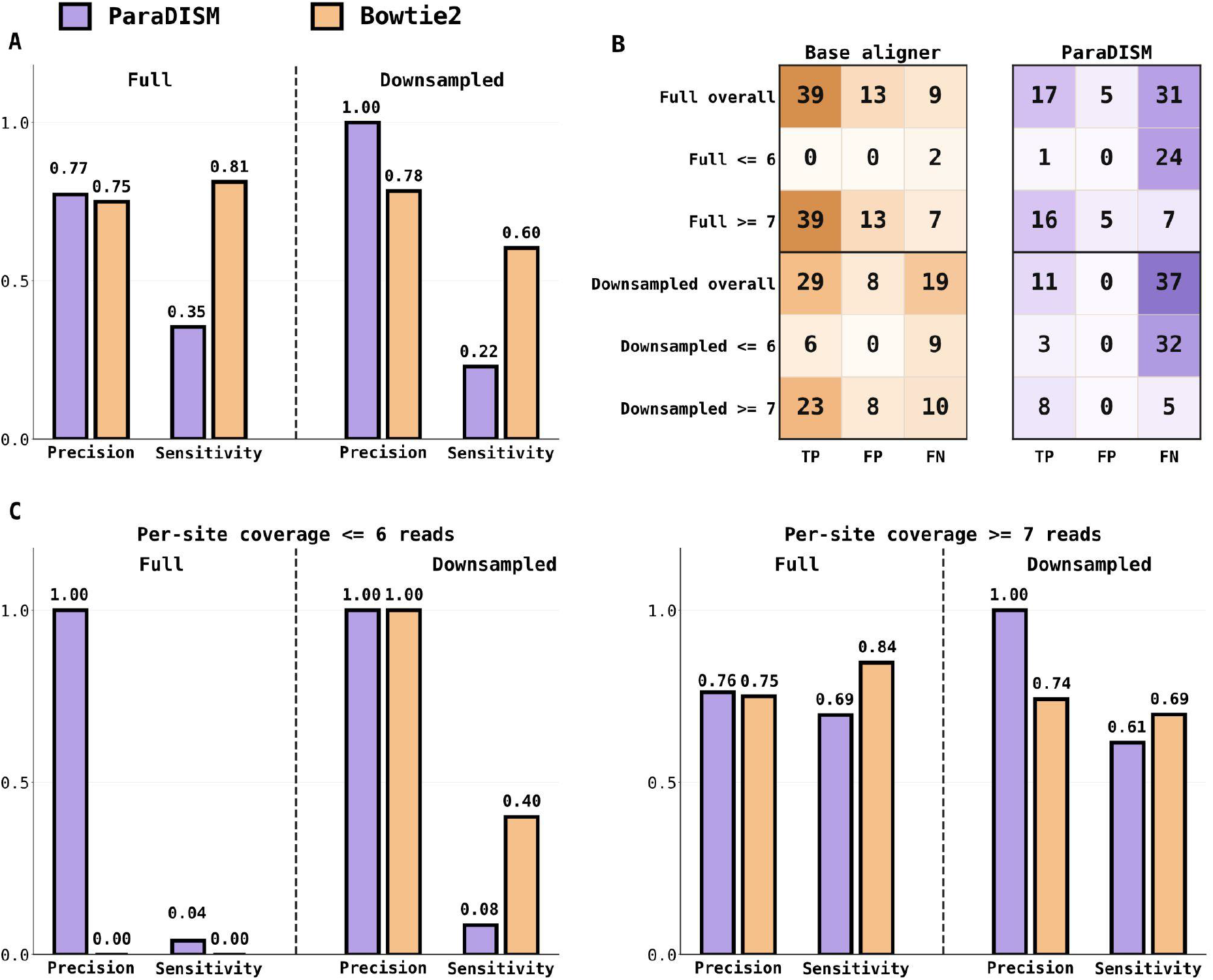
Variant calling performance on the GIAB HG002 PKD1 benchmark. (A) Overall precision and sensitivity of SNV calls obtained from ParaDISM versus the base aligner (Bowtie2) on the full and downsampled (∼10x) GIAB HG002 short-read dataset. (B) SNV call count matrix for the base aligner vs ParDISM. Rows indicate dataset and coverage stratum, and columns indicate the numbers of true positives (TP), false positives (FP), and false negatives. (C) Precision and sensitivity stratified by coverage, using a threshold of 6 reads to separate low-coverage and higher-coverage sites in the full and downsampled datasets.

On the full dataset, *ParaDISM* resulted in five false positives and 17 true positives compared to 13 false positives and 39 true positives of Bowtie2, corresponding to a moderately increased precision from 0.75 to 0.77. The moderate increase in precision was caused by the fact that *ParaDISM* is limited by the reference sequence homology rather than coverage, while Bowtie2 is not. Accordingly, the increased coverage allowed Bowtie2 to detect ten more true SNVs (at the cost of reporting five more false SNVs), while *ParaDISM* detected only six more SNVs (at the cost of reporting also five more false SNVs).

The lower sensitivity of SNVs called on *ParaDISM* alignments compared to Bowtie2 was a direct consequence of a more stringent approach to read placement. Accordingly, in the 10x downsampled data set, the false negative calls of *ParaDISM* were located predominantly in the regions of low coverage, with 32 out of 37 false negative calls located in sites covered with less than 7 reads and only 5 false negative calls located in sites covered with at least 7 reads (Fig. 3). Therefore, the overall decrease in sensitivity from 0.60 to 0.23 is easily manageable in practice. The BAM files produced by *ParaDISM* can be inspected manually, and regions with low coverage can instantly be spotted. Based on other SNVs called by the pipeline, the user may decide whether the low coverage regions warrant further investigation with other approaches such as PCR. On the other hand, detecting and confirming false positives produced by the standard pipelines is a laborious process with no clear guidelines.

### *GNAQ* case study: *ParaDISM* mitigates misalignment artifacts

To illustrate how *ParaDISM* can protect against misalignment artifacts, we applied it to re-analyze SNVs in *GNAQ* and *GNAQP1* sequences in five tumour WES data sets in which previous reports suggested the possibility of false SNVs caused by misalignment artifacts [6]. In the SNVs called by FreeBayes from *ParaDISM* alignments, no alternative base on *GNAQ* was observed at either hotspot position originally reported by Li et al. (2019) across the cohort (*GNAQ*:206083 ALT=0/200; *GNAQ*:206100 ALT=0/284; Table 1). At the three *GNAQP1* SNP positions described by Lim et al. (2022), the full T,C,T ALT triplet was not observed: the rs3730150 ALT base (T) was present at low frequency at *GNAQP1*:765 (13/153 reads), whereas the rs3730148 ALT base (C) at *GNAQP1*:786 and the rs3730153 ALT base (T) at *GNAQP1*:826 were not observed (0/162 and 0/237 reads, respectively; Table 1).

**Table 1:** ALT evidence at GNAQ hotspot sites and linked GNAQP1 SNP sites in ParaDISM gene-specific alignments. Per-base evidence at two GNAQ hotspot positions (GNAQ:206083, GNAQ:206100) and at the three linked GNAQP1 SNP positions reported in Lim et al. (2022) (GNAQP1:765, 786, 826), summarised from ParaDISM sequence-specific BAM outputs for five tumour exomes. For GNAQ sites, ALT denotes any base other than the only observed base at that position. For GNAQP1 sites, counts are shown for the Lim et al. ALT bases (T,C,T). DP denotes read depth.

| Sample | GNAQ:206083<br>ALT/DP | GNAQ:206100<br>ALT/DP | GNAQP1:765<br>T/DP | GNAQP1:786<br>C/DP | GNAQP1:826<br>T/DP |
| --- | --- | --- | --- | --- | --- |
| SRR5602384 | 0/28 | 0/35 | 6/6 | 0/0 | 0/1 |
| SRR5602389 | 0/51 | 0/69 | 4/54 | 0/62 | 0/86 |
| SRR5602393 | 0/47 | 0/69 | 1/25 | 0/25 | 0/32 |
| SRR5602414 | 0/43 | 0/69 | 2/36 | 0/37 | 0/52 |
| SRR5602419 | 0/31 | 0/42 | 0/32 | 0/38 | 0/66 |
| Total | 0/200 | 0/284 | 13/153 | 0/162 | 0/237 |

These results illustrate how *ParaDISM* is less prone to align reads in ambiguous cases compared to standard aligners. In this case study, the pipeline did not provide a conclusive answer as to whether the hotspots are genuine variants or misalignment artifacts, indicating that this question may warrant further studies. This way, *ParaDISM* avoids the “confident uncertainty” that may affect the standard aligners and cause them to output misaligned reads with a high MAPQ score. The latter can happen especially when, due to the presence of variants or gene conversion, the incorrect location has a higher alignment score than the correct one. Since *ParaDISM* does not rely on alignment score alone, its results are less affected by a potentially misleadingly high MAPQ.

### ADPKD case study: *ParaDISM* detects medically relevant variants

In the final case study, we demonstrate that, despite the lower sensitivity, *ParaDISM* can identify medically relevant variants. We applied *ParaDISM* to analyze srHTS data of patients diagnosed with ADPKD (three panel, 15 WES datasets) and compared it to the variant outputs of a more standard approach that is not paralog-aware.

The NextGENe v.2.4.2.3 (SoftGenetics, USA) pipeline reported potentially causative variants in 13 of the 15 ADPKD families: a splicing mutation in *PKD2* and 14 shortlisted variants in *PKD1* which included 5 truncating SNVs, 8 missense variants and 1 in-frame 3bp deletion. No variants were shortlisted in 2 families.

Analysis of the same datasets using *ParaDISM* confirmed 5 truncating variants, 5 missense variants and the in-frame deletion variant. Furthermore, *ParaDISM* did not call 2 missense variants reported by the NextGENE pipeline (p.C508R in exon 7 and p.F3873V in exon 42) which alternative sequencing methods had shown to be absent from *PKD1* despite prior shortlisting from srHTS datasets, and were likely caused by misalignment artifacts. Finally, *PKD1*p.R739Q, a commonly reported benign variant which had been called in a number of patient datasets, was not called by *ParaDISM*. This variant is currently labelled as a low quality variant in the latest gnomAD database (v4.1.0). On the other hand, a potentially causative missense variant confirmed to be on exon 11 of *PKD1* by long-range PCR and Sanger sequencing of nested PCR product was not detected by *ParaDISM*. This variant was located in a region with a low coverage of *ParaDISM* alignments. The reads bearing this variant were identified in the BAM file containing reads that did not pass the paralog disambiguating conditions.

## Conclusions

Highly similar sequences in the genome are a major challenge for srHTS technologies. Standard bioinformatic pipelines that may force an alignment in these regions can result in misplaced reads. These misalignment artifacts can propagate through bioinformatic pipelines and result in false positive and false negative variant calls. Both types of errors have equally important implications for clinical practice and biomedical research. False positive calls happen when a called variant is not present in the genome, and can result in patients being subjected to unnecessary treatments. False negative calls happen when a software fails to detect a variant present in the genome and can result in misdiagnosing affected patients as healthy. Computational tools cannot avoid both types of false results at once, resulting in a trade-off where increasing the sensitivity causes decreased specificity and *vice versa*.

*ParaDISM* is optimized for precision and explicitly reports unresolved ambiguity rather than masking it as confident alignment. This approach is appropriate for those diagnostic settings in which false positives can be more harmful than reduced sensitivity. Furthermore, the rate of false negative calls can be mitigated in practice by inspecting the coverage of *ParaDISM* alignments. The pipeline achieves a higher precision compared to standard aligners by using multiple sequence alignment comparison of reference sequences and aligning only reads with unambiguous evidence for their sequence of origin. As demonstrated by the case studies, *ParaDISM* can be used with different types of srHTS data: panel sequencing (e.g. the *PKD1* case studies), whole-exome sequencing (e.g. the *GNAQ* and *PKD1* case studies) and whole-genome sequencing (e.g. the GIAB benchmark). While the pipeline requires users to provide the specific sequences of interest and can only analyze a single set of reference sequences at a time, a single srHTS data set can be reused to study multiple genes by running *ParaDISM* with different sets of reference sequences.

Future work can further improve the performance of *ParaDISM*. Currently, while the software tackles disambiguation between duplicated genes, it does not explicitly disambiguate alternative placements within duplicated regions of a single gene. Instead, it evaluates the alignment records reported in the input SAM and assigns a read pair only when those records provide unique support for exactly one gene. Furthermore, if a patient has variants in the “anchoring” sites, especially resulting from gene conversion and thus matching other sequences, *ParaDISM* may not be able to place reads around them. Finally, *ParaDISM* may potentially produce false negative calls in heterozygotic mutations if reads from only one haplotype can be placed confidently. Cross-checking with other bioinformatic pipelines or inspecting the unassigned reads is recommended to detect potential SNVs in reads that did not pass the paralog disambiguating criteria.

## Availability and requirements

**Project name:** *ParaDISM*

**Project home page:** github.com/BioGeMT/ParaDISM

**Archived version:** https://zenodo.org/uploads/18493364

**Operating system(s):** Platform independent **Programming language:** Python 3

**Other requirements:** Python (>=3.9); MAFFT; Bowtie2 and/or BWA-MEM2 and/or minimap2; FreeBayes; SAMtools

**License:** MIT

**Any restrictions to use by non-academics:** None

## List of abbreviations

ADPKD: autosomal dominant polycystic kidney disease
ALT: alternate allele
AF: allele frequency
BAM: Binary Alignment/Map
DP: read depth
FN: false negative
FP: false positive
GIAB: Genome in a Bottle
GRCh37: Genome Reference Consortium Human Build 37
GRCh38: Genome Reference Consortium Human Build 38
HTS: high-throughput sequencing
NCBI: National Center for Biotechnology Information
NGS: next-generation sequencing
NIST: National Institute of Standards and Technology
NKTCL: natural killer/T-cell lymphoma
MSA: multiple sequence alignment
ParaDISM: Paralog Disambiguating Mapper
PCR: Polymerase Chain Reaction
QUAL: variant quality (QUAL score)
SNP: single-nucleotide polymorphism
SNV: single-nucleotide variant
SRA: Sequence Read Archive
srHTS: short-read high-throughput sequencing
TP: true positive
UNMET: UNMET score (exome-wide short-read sequencing difficulty metric)
VCF: Variant Call Format

## Declarations

### Ethics approval and consent to participate

The ADPKD study was approved by the University of Malta Research Ethics Committee (ethics approval number FHS031/2014). Written informed consent was obtained from all participants.

### Consent for publication

Not applicable.

### Availability of data and materials

The datasets generated and/or analysed during the current study are available in the Zenodo repository, https://zenodo.org/uploads/18493364. The Zenodo record contains the outputs generated in this study (including simulation summary tables and figure source data, GIAB benchmarking *ParaDISM* outputs and variant calls, and *GNAQ/GNAQP1 ParaDISM* outputs and variant calls). The original/raw clinical sequencing reads (whole-exome/panel FASTQ files) for the ADPKD cohort and their *ParaDISM* outputs are available upon reasonable request as per ethics approval for this data. For the *PKD1/PKD1P1-6* simulations, the Zenodo record includes the summary tables used to generate Fig. 2 (overall precision/sensitivity/specificity and aggregate read assignment matrices reported as mean ± standard deviation across 1,000 seeds). Simulation configuration (DWGSIM parameters, reference FASTA, and the list of random seeds) and scripts to regenerate per-seed outputs are available at github.com/BioGeMT/ParaDISM. Due to size, the complete set of per-seed raw simulation outputs (e.g., FASTQ/BAM/) for all 1,000 runs was not archived.

Public input datasets analysed during the study are available from Genome in a Bottle (GIAB) for HG002: GIAB HG002 v4.2.1 (GRCh38) benchmark VCF: https://ftp-trace.ncbi.nlm.nih.gov/ReferenceSamples/giab/release/AshkenazimTrio/HG002_NA24385_son/NISTv4.2.1/GRCh38/HG002_GRCh38_1_22_v4.2.1_benchmark.vcf.gz (ftp-trace.ncbi.nlm.nih.gov), GIAB HG002 v4.2.1 (GRCh38) benchmark regions BED: https://ftp-trace.ncbi.nlm.nih.gov/ReferenceSamples/giab/release/AshkenazimTrio/HG002_NA24385_son/NISTv4.2.1/GRCh38/HG002_GRCh38_1_22_v4.2.1_benchmark_noinconsistent.bed (ftp-trace.ncbi.nlm.nih.gov), GIAB HG002 Illumina 2×250 FASTQs (reads directory): https://ftp-trace.ncbi.nlm.nih.gov/ReferenceSamples/giab/data/AshkenazimTrio/HG002_NA24385_son/NIST_Illumina_2x250bps/reads/.

Public datasets from NCBI SRA for the five tumour exome datasets used in the *GNAQ/GNAQP1* case study, NCBI SRA study (SRP107053): https://www.ncbi.nlm.nih.gov/sra/?term=SRP107053 (contains SRR5602384, SRR5602389, SRR5602393, SRR5602414, SRR5602419). All scripts and configuration required to reproduce the analyses and regenerate the figures are available in the *ParaDISM* GitHub repository (github.com/BioGeMT/ParaDISM) and are archived in the Zenodo record.

### Authors’ contributions

RF, MC and PA conceived the study and SBW contributed to the development of the idea. DT implemented the pipeline and ran the computational experiments. AB supervised the implementation. MC and RF interpreted the results. PA advised on the implementation of the pipeline and interpretation of the results. FBC, MCM and JZ carried out the initial analysis of ADPKD cases and wet-lab validations supervised by RF. SBW coordinates the Malta NGS project which includes the ADPKD study. MC coordinated the work. MC, DT, RF and AB wrote the manuscript. All authors have read and critically revised the manuscript and approved the final version.

